# How sex shapes transcriptome evolution in the songbird brain

**DOI:** 10.1101/2025.08.21.671601

**Authors:** Isaac Miller-Crews, Sara E. Lipshutz, Ben Fulton, Jason Bertram, Matthew W. Hahn, Kimberly A. Rosvall

## Abstract

Sex differences have long captivated scientists, yet the evolutionary rate of change in sex-biased gene expression has not been directly quantified. To address this issue, we introduce new options in CAGEE (Computational Analysis of Gene Expression Evolution), specifically unbounded Brownian motion and variable evolutionary rates among genes. We applied these features to brain transcriptomes of ten songbird species, half of which convergently evolved obligate cavity-nesting, an element of reproductive ecology linked to sex-specific changes in competitive behavior. We find that the degree of sex-bias – measured as male:female expression ratio for each gene – evolves twice as fast on the Z chromosome versus autosomes, but Z gene expression does not evolve at different rates among sexes. Most Z-linked genes are male-biased in their expression, but not all. These sex-balanced genes are not skewed in their rate of evolution, contrary to the hypothesis that some genes experience selection for balance and consequently evolve more slowly. Finally, the degree of sex-bias in gene expression evolves more quickly along obligate-cavity nesting lineages, suggesting that sex-specific selection may shape the evolution of brain sex differences, or lack thereof. Together, these results provide new insights into the interplay between sex and gene expression evolution.

## INTRODUCTION

Biologists have long been captivated by why species vary in the degree of sexually differentiated traits (Darwin, 1871). This variation may come through various mechanisms at different levels of biological organization, including evolutionary changes to sex chromosomes, gene expression, and protein sequences (Rowe et al., 2018). For example, within many different species, we know that sex differences in brain gene expression correspond to sex differences in other components of the phenotype (Harris & Hofmann, 2014; Renn et al., 2008) (but see (De Vries, 2004). Comparative studies of small numbers of lineages further re-iterate connections between sex differences in gene expression and trait evolution (Houle & Cheng, 2021; Ko et al., 2021; Schumer et al., 2011). There is also evidence that male-vs. female-biased genes differ in how they evolve (Mank, 2017; Rowe et al., 2018), yet the evolutionary rate of those sex differences in gene expression has not been directly quantified, underscoring the need for empirical testing (Tosto et al., 2023).

One challenge to filling this knowledge gap is methodological. Most phylogenetic analyses of transcriptomic data estimate the rate of evolution in each gene individually, using post-hoc analyses to infer larger patterns across the transcriptome (e.g. (Pease et al., 2022; Yang et al., 2019)). Recently, the software CAGEE (Computational Analysis of Gene Expression Evolution) was developed to assess evolutionary rates while accounting for the fact that genes evolving along a lineage do not evolve independently from one another (Bertram et al., 2023). CAGEE v1.0 estimates rates of expression evolution across an entire transcriptome or a subset of the transcriptome (e.g. a particular chromosome). Variation in these rates can be modeled across one or more branches of a phylogeny (Fig. 1A), or across different tissues, treatments, or sampling timepoints within multiple species. However, CAGEE v1.0 and later v1.1 (which fixed several bugs) otherwise assume all genes share the same average rate of evolution. This assumption could obscure interesting patterns, considering that different genes could evolve at different rates. The existing CAGEE software also uses a bounded Brownian motion model, which only considers continuous positive traits (e.g. absolute gene expression levels). While this approach works for some comparisons, for instance analyses of each sex independently, the rate of evolution of sex bias is better captured as a male-to-female ratio of gene expression (hereafter, ‘M:F ratio’). Once log-transformed for analyses, this metric includes negative values, thereby necessitating an unbounded Brownian motion model. By expanding CAGEE to include an unbounded Brownian motion model (Fig. 1B), alongside the option for different evolutionary rates among subsets of genes (Fig. 1C), we can directly test long-standing predictions related to the evolution of sex-biased genes, and in doing so, we also broaden the types of research questions that can be addressed in comparative transcriptomics.

**Figure 1:**
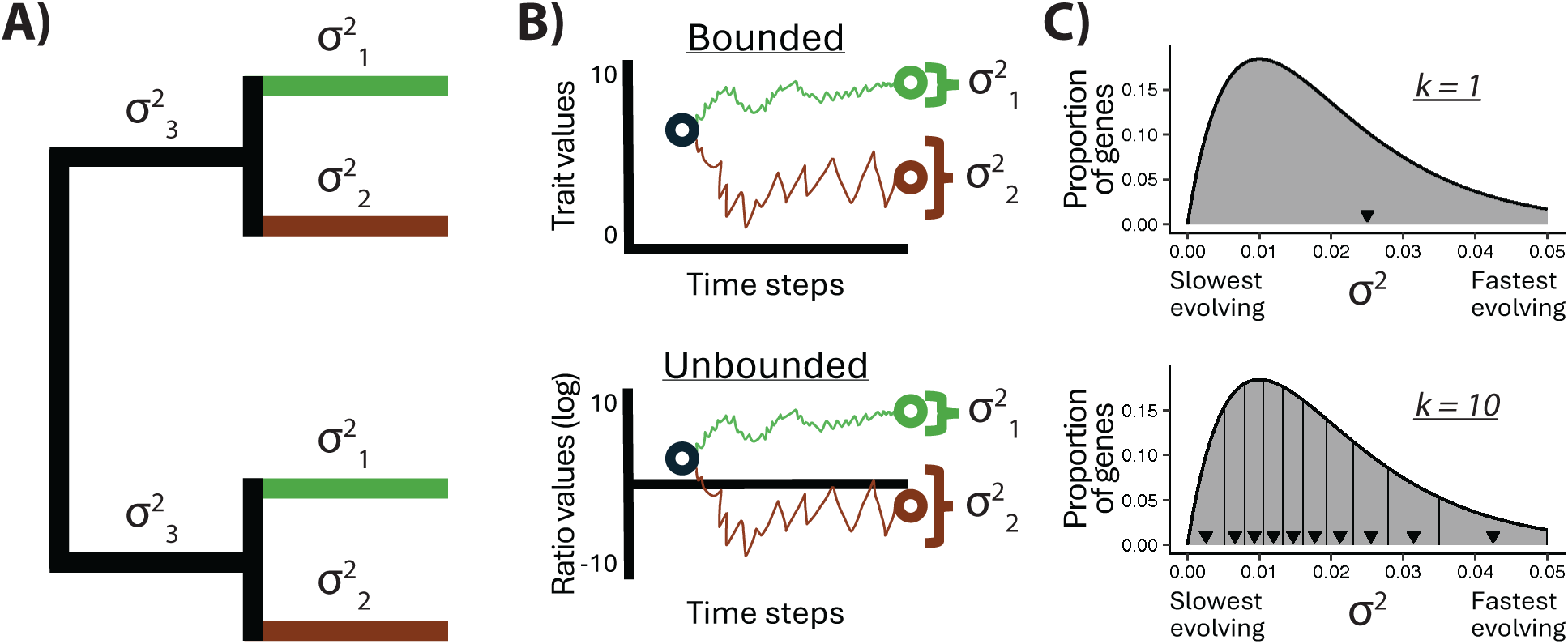
CAGEE v1.2 now integrates two new features to extend the original CAGEE. **A)** With CAGEE v1.0, a transcriptome-wide evolutionary rate (σ^2^) can be calculated across the phylogeny, or different σ^2^-values can be calculated for specific branches (green vs brown lineages vs black internal branches). σ^2^ represents the evolutionary variance over time, with larger values indicating a faster amount of change. CAGEE also uses trait values at the tips of the tree to conduct ancestral state reconstruction at internal nodes. **B)** CAGEE v1.0 models incremental changes over time with a bounded Brownian motion model (top), for use with continuous positive traits (e.g. gene expression). CAGEE v1.2 also includes an unbounded Brownian motion (bottom), for use with any continuous trait (e.g. ratio of gene expression between sexes, converted to a log scale). **C)** CAGEE v1.0 calculates a single σ^2^ for the average σ^2^ (black triangle) across the distribution of all genes (top). CAGEE v1.2 allows for variation in σ^2^ among genes (bottom) by fitting data to a discrete-gamma distribution, calculating *k* equiprobable rate categories (e.g. k = 10), and assigning the average rate (triangle) to all genes in that category.

Songbirds are well-suited to these questions for several reasons. For one, avian chromosomes are relatively stable, with a high degree of synteny (Ellegren, 2010; Shetty et al., 1999), including on the Z sex chromosome (Nanda et al., 2008; Zhou et al., 2014). Z genes tend to evolve more rapidly than those of autosomes, in both coding sequence (Wright, Harrison, et al., 2015) and expression (Dean et al., 2015), and sex differences in expression may be an important driver of behavioral variation, even controlling for hormones or gonadal tissue (Wade et al., 1999). Gene expression on the Z is also 30-40% higher in males compared to females (Itoh et al., 2007; Julien et al., 2012). This is a notable feature because male birds have two Z chromosomes and females have one Z and one W. Most animals handle the problem of chromosome number variation between the sexes via dosage compensation, in which expression outputs are equalized between the shared sex chromosome in the homogametic and heterogametic sexes (Chen et al., 2020; Gu & Walters, 2017). However, birds do not exhibit global dosage compensation (Graves, 2016); instead, individual Z genes vary in their degree of sex difference, and some genes exhibit relative balance in expression between the sexes (Itoh et al., 2010; Ko et al., 2021; Naurin et al., 2011; Wolf & Bryk, 2011), prompting questions as to whether these Z genes are under selection to be balanced. If so, we may expect that sex-balanced genes would maintain balance across the phylogeny, and therefore, these more balanced genes would exhibit both a balanced ancestral state and a slower evolutionary rate in their degree of sex bias. Whether some Z genes consistently exhibit balance due to evolutionary constraint is unclear and requires analysis of the degree of sex bias.

Beyond these chromosomal considerations, there also may be ecological drivers of gene expression evolution, for example with foraging guilds (Nesta & Ledon-Rettig, 2025), subterranean living (Partha et al., 2017), predation pressure (Ghalambor et al., 2015), or sexual selection (Harrison et al., 2015). For birds, cavity-nesting is another relevant ecological factor, which has arisen independently over 30 times (Fang et al., 2018). Obligate secondary cavity-nesters require a hole in a tree or other substrate in order to reproduce, and they cannot excavate one for themselves. Cavity-nesting is also associated with distinct selection pressures and phenotypic effects, including slower developmental trajectories, altered predation and thermal regimes, and more competition for limited breeding sites (Bosque & Bosque, 1995; Drury et al., 2020; Martin et al., 2017; Martin & Li, 1992; Zenil-Ferguson et al., 2023). Critically for our consideration of sex and its role in evolution, obligate cavity-nesting is associated with female-specific increases in conspecific aggressiveness (Lipshutz et al., 2025; Lipshutz & Rosvall, 2021). This increase in aggression is thought to stem from heightened pressures on cavity-nesting females to find and secure a limited nesting site – pressures that are softened among females in more flexibly nesting species whose reproductive success does not require such a restricted ecological niche. How these ecological changes relate to the pace of sex-associated gene expression evolution is unknown.

Here, we use brain gene expression data from 10 songbird species to explore how sex differences in gene expression evolve, using a new version of CAGEE (v1.2). This updated CAGEE introduces two new features: (i) an unbounded Brownian motion model and (ii) the option to allow genes to have different evolutionary rates via gamma-distributed rate categories (see Fig. 1). We apply these new features to test three hypotheses: First, we hypothesize that the degree of sex differences in gene expression on the Z will evolve more quickly than that of the autosomes, driven by faster evolution of the female Z. Second, we hypothesize that some Z genes may be selected for more balanced expression between the sexes; if so, there will be an overrepresentation of Z genes with both more balanced expression and a slower evolutionary rate for the M:F ratio. Third, we hypothesize that changes in sex-specific selection will shape how sex differences in the brain evolve, with sex-biased gene expression evolving more rapidly alongside the transition to an obligate cavity-nesting phenotype. Together, these hypotheses, and the new CAGEE features we test alongside them, open new doors into our understanding of sex and its interaction with transcriptomic evolution.

## MATERIALS AND METHODS

### Sample collection

Samples were collected from mid-March to mid-May 2018-2021 in Indiana, Kentucky, or Illinois, USA, as detailed in Lipshutz et al. (2025). This earlier paper controlled for sex in an analysis of convergent evolution of gene expression and behavior; here, we develop and apply novel analyses that dive deeper into the degree of sex differences and their pace of evolutionary change. We focused on 10 songbird species, initially selected for a pairwise comparison of obligate secondary cavity nesting (listed first) vs. their more flexibly nesting relatives (listed second): Eurasian tree sparrow (*Passer montanus*) vs. house sparrow (*Passer domesticus*), prothonotary warbler (*Protonotaria citrea*) vs. yellow warbler (*Setophaga petechia*), Eastern bluebird (*Sialia sialis*) vs. American robin (*Turdus migratorius*), house wren (*Troglodytes aedon*) vs. Carolina wren (*Thryothorus ludovicianus*), tree swallow (*Tachycineta bicolor*) vs. barn swallow (*Hirundo rustica*) (Fig. 2A). All birds were in the same pre-breeding stage of territory establishment and pair bonding, confirmed via behavioral observations and post-mortem gonadal inspection.

**Figure 2:**
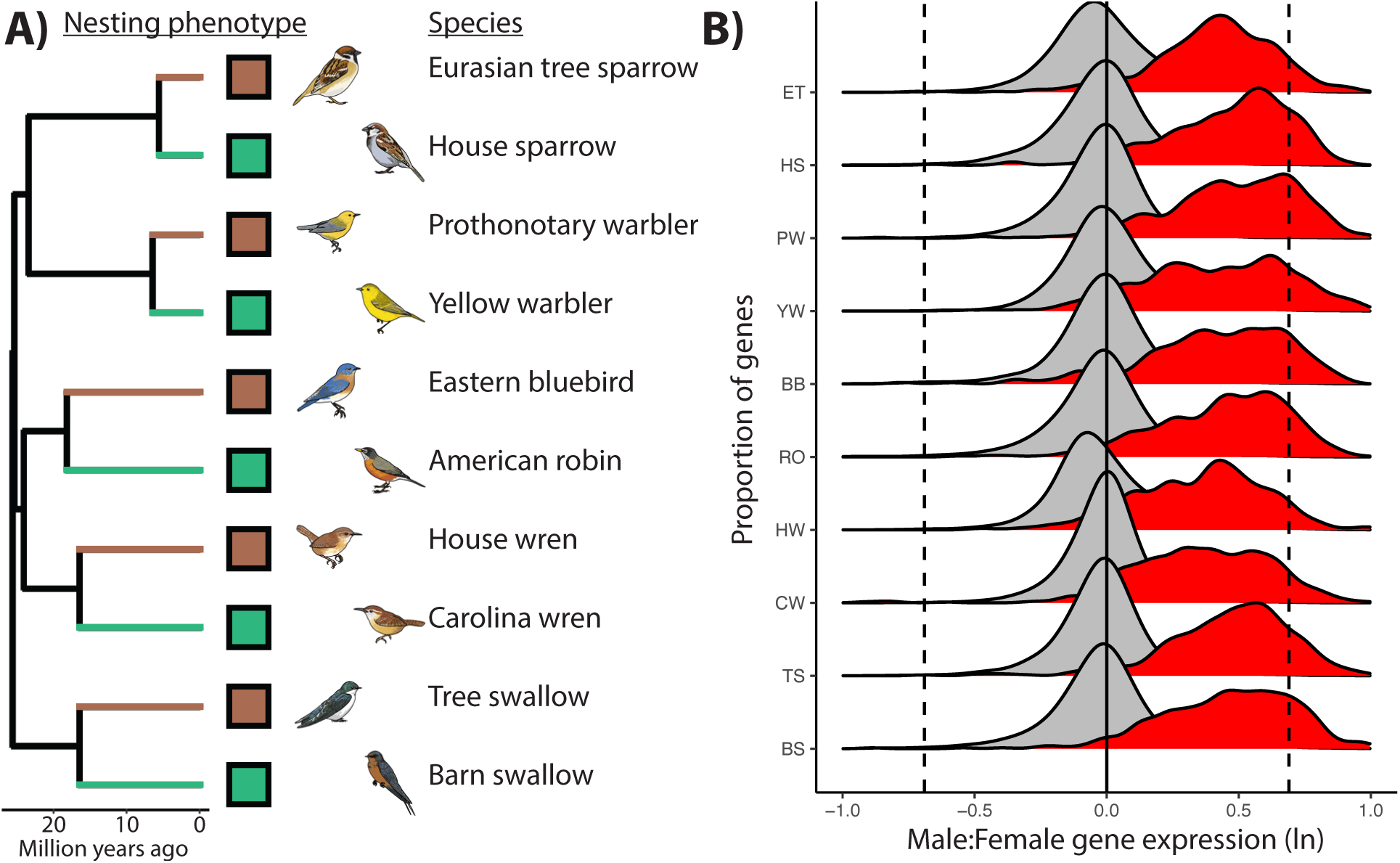
**A)** Phylogeny of 10 focal species from TimeTree (Kumar et al., 2022). External branches are colored by cavity-nesting phenotype (brown = obligate cavity-nesting, green = not obligate cavity-nesting), with internal branches colored black (*illustrations by Tessa Patton*). **B)** Density plots show M:F gene expression ratio for each species. Each species is a row, with autosomal genes in grey and Z chromosome genes in red. Positive values indicate higher expression in males compared to females, and vice versa.

Each subject was captured on its territory and euthanized with an overdose of isoflurane followed by immediate decapitation, brain dissection, and flash freezing on dry ice (n = 6/sex per species, with n=7 for female tree swallow, for total n=121). Brains were later microdissected following (Soma et al., 2003), and we focused on the ventromedial telencephalon (VmT), which includes the avian medial amygdala (or TnA (Reiner et al., 2004) or AMV (Mello et al., 2019)), bed nucleus of the stria terminalis (BNST), and lateral septum (LS) because these regions play important roles in regulating and responding to aggression across many species, including cavity-nesting birds (Bentz et al., 2019; Horton et al., 2014; O’Connell & Hofmann, 2011). We isolated total RNA using Trizol (Invitrogen, USA).

### Sequencing, alignment, and mapping

Sequencing and pre-processing is detailed in Lipshutz et al. (2025); briefly, we performed RNAseq (Illumina Nextseq500, 75 cycle), which yielded ∼28 million reads per sample. Transcriptomes for each species were assembled with Trinity version 2.13.2 (Grabherr et al., 2011) and aligned to the zebra finch (Taeniopygia guttata) proteome (GCF_003957565.2_bTaeGut1.4.pri_protein.faa) (Rhie et al., 2021), yielding 10,672 orthologous genes present across all 10 species (for details on ortholog identification see Lipshutz et al., 2025)).

We assigned chromosomal location to genes using Ensembl (Harrison et al., 2024) and the zebra finch reference genome. Because this reference is from a male that does not have a W chromosome, we did not assign W-linked genes. W-linked genes tend to experience purifying selection (Bellott et al., 2017; Moghadam et al., 2012), and they may be involved in restoring dosage balance with their Z-linked homologs (Smeds et al., 2015; Xu & Zhou, 2020). Nonetheless, they are relatively small in number (∼30-50 genes (Ko et al., 2021)), such that their absence in our dataset should have a minimal impact on our broader findings.

### Modelling the evolution of gene expression in CAGEE

Using normalized gene expression counts from DESeq2 (Supplemental Dataset S1), we calculate three main dependent variables: (1) median gene expression for each species; (2) median gene expression for males and females separately, within each species; and (3) ratio of male median and female median gene expression within a species (M:F ratio), which provides insight into the degree of species-level sex differences for each gene (Supplemental Dataset S2). This lattermost metric is the first implementation of an unbounded Brownian motion model, now available in CAGEE v1.2 (https://github.com/hahnlab/CAGEE; Fig. 1B). In practice, both the original bounded Brownian and the new unbounded Brownian motion models require an artificial upper bound to allow numerical analyses to finish in a reasonable time; the new unbounded Brownian motion model also includes an artificial lower bound that is simply the negative reflection of the upper bound.

### Modelling the role of sex and sex chromosomes

To test our first main hypothesis, we focus on the role of the Z chromosome in the rate of gene expression evolution, using both pre-existing and new features of CAGEE (see Introduction). We apply a series of nested models to calculate the average rate evolution per million years (σ^2^) and a log-likelihood (-lnL) that is used to compare fit among iterations of competing models (Bertram et al., 2023), applied separately to each of the three dependent variables described in the previous paragraph. We could then determine the best fitting model among the nested models with the lowest-lnL and the fewest number of parameters. Specifically, iterations of Model 1 ignore sex, asking only whether the species median rate of gene expression evolution is faster on the Z vs. autosomes (Table 1). Iterations of Model 2 isolate both sex and sex chromosome, asking whether median male and median female gene expression levels evolve at different rates, and whether any differences are driven by the Z chromosome in one or both sexes (Table 2). Iterations of Model 3 focus on the M:F ratio of gene expression, asking directly in a single phylogenetic model how the degree of sex bias evolves on the Z vs. autosome (Table 3).

**Table 1.** Species gene expression evolutionary rate.

| <u>Model</u> | <u>-lnl</u> | <u>rate</u> | <u>$\sigma^2$</u> |
| --- | --- | --- | --- |
| All | 337733 | All | 0.01138 |
| **Autosomal vs Z** | **337690** | **A** | **0.01128** |
|  |  | **Z** | **0.01420** |

**Table 2.**
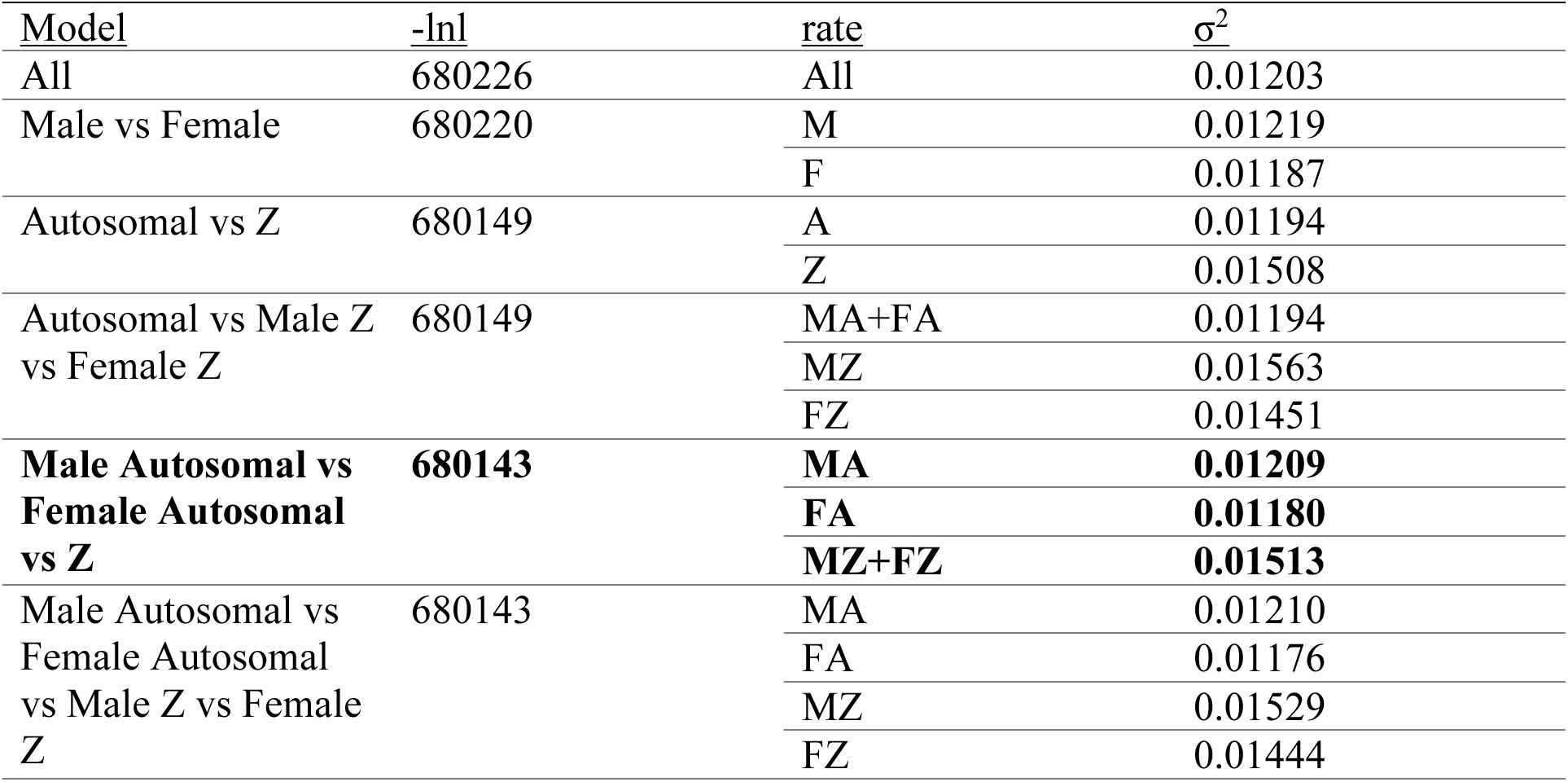
Gene expression evolutionary rate per sex.

**Table 3.**
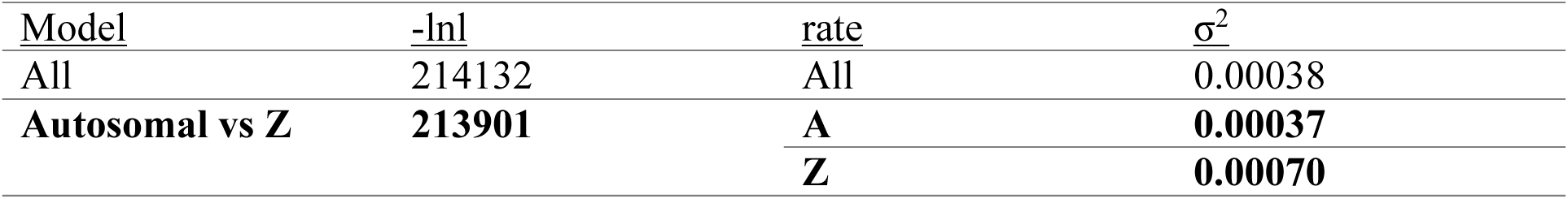
M:F ratio gene expression evolutionary rate.

### Modelling with distinct rate categories

We also use the second new feature in CAGEE v1.2—gamma-distributed rate categories—to explore how the rate of evolution differs among genes on the same branch of the phylogeny (Fig. 1C). This feature allows multiple, discrete equiprobable rate categories drawn from a gamma distribution calculated from the data. We assume that the shape parameter (a) is equal to the scale parameter (b), finding the best-fit model to the whole dataset with the specified *k* (see Mendes et al., 2020; Yang, 1996). It is possible for genes to be partial members of multiple rate categories, in that the rate would better be reflected by the inclusion of an additional intermediate category, but a gene can also be assigned to its most likely rate category using maximum likelihood; the σ^2^ for that gene is then presumed to be the σ^2^ for that category. After validating our approach in a simulated dataset (see Supplement §2, Fig. S3), we modeled our ∼10k gene dataset with *k*=10 rate categories, for both species (median) gene expression (dependent variable 1) and M:F ratio (dependent variable 3). We chose *k*=10 rate categories to increase granularity while maintaining a robust number of genes in each category, targeting on average 1,000 genes per category. Although it is not possible to identify an optimal value of *k* for a dataset (Yang 1996; Mendes et al. 2020), our validations across various *k* values demonstrate that this choice does not affect our conclusions (see Supplement §2, Fig. S4).

Outputs from these variable rate models were used in several downstream analyses, assigning individual genes to a single rate-category. A) We used a one-tailed Fishers exact test to ask whether genes whose expression evolves quickly are also genes whose sex-ratio of expression evolves quickly (and vice versa, for slowly evolving genes). B) We used a one sample two-tailed t-test to ask directly whether, in each rate category, there was a different percentage of Z chromosome genes compared to other chromosomes, for both gene expression and M:F ratio. C) We used gene ontology (GO) analyses to ask whether genes with the fastest (or slowest) rates of evolution are enriched for specific functional pathways, for both species (median) gene expression and M:F ratio (elaborated in Supplement §4).

### Modelling ancestral states of male:female ratio

In accordance with our second main hypothesis focused on the Z, we used an ordinal logistic regression to ask whether ancestral state of sex bias (i.e. M:F ratio at the root node of the tree (Bertram et al., 2023)), predicts the evolutionary rate category, as is expected if there is selection for some genes to be more balanced in their expression between the two sexes.

### Modelling the role of nesting strategy

Finally, for our third main hypothesis, we focus on cavity nesting as a potential ecological driver of sex-specific gene expression evolution. We modeled species (median) gene expression (dependent variable 1) with bounded Brownian motion (Table 4), and M:F ratio (dependent variable 3) with unbounded Brownian motion (Table 5). In each case, we compared three model iterations: i) all branches together (i.e. one σ^2^ across the phylogeny); ii) internal and external branches having separate values of σ^2^; iii) and internal, obligate cavity-nesting, and more flexibly nesting branches each having their own σ^2^ parameter. To avoid over-parameterization, these models were agnostic to chromosome and used a single (average) σ^2^ for all genes.

**Table 4.** Cavity nesting gene expression evolutionary rate.

| <u>Model</u> | <u>-lnl</u> | <u>rate</u> | <u>$\sigma^2$</u> |
| --- | --- | --- | --- |
| All | 337733 | All | 0.01138 |
| Internal vs External | 336583 | Internal | 0.00629 |
|  |  | External | 0.01360 |
| **Nesting Phenotype** | **336577** | **Internal** | **0.00625** |
|  |  | **Obligate** | **0.01328** |
|  |  | **Flexible** | **0.01401** |

**Table 5.** M:F ratio cavity nesting gene expression evolutionary rate.

| **Table 5. M:F ratio cavity nesting gene expression evolutionary rate** |  |  |  |
| --- | --- | --- | --- |
| <u>Model</u> | <u>-lnl</u> | <u>rate</u> | <u>$\sigma^2$</u> |
| All | 214132 | All | 0.00038 |
| Internal vs External | 208787 | Internal | 0.00006 |
|  |  | External | 0.00055 |
| **Nesting Phenotype** | **208723** | **Internal** | **0.00006** |
|  |  | **Obligate** | **0.00059** |
|  |  | **Flexible** | **0.00050** |

## RESULTS

### Differential expression analysis of sex bias per species

Most of the genes that differ in their expression between female and males were found on the Z (see Supplement §1, Fig. S1). Z genes have a mainly male-biased distribution (i.e. higher in males than in females), but a small fraction of Z genes show approximately equal (balanced) or female-biased expression (see Fig. 2B), providing a strong foundation for testing hypotheses on the degree of sex bias in gene expression.

### Modelling the role of sex and sex chromosomes

Focusing first on median gene expression and without considering sex, we found that rates of evolution on the Z chromosome are faster than on the autosomes (σ^2^ = 0.01420, vs. 0.01128); this model allowing for separate rates among chromosomes is a significantly better fit than the model that does not allow for different rates (Table 1). In models that directly include both sex and sex chromosome (i.e. testing our first main hypothesis), the best fit model allows for three rates of gene expression evolution: one for autosomes in males (σ^2^ = 0.01219), another (slightly lower) value for autosomes in females (σ^2^ = 0.01187), and a third (higher) value for the Z, regardless of sex (σ^2^ = 0.01513; Table 2). To test the robustness of these results, we used a random sub-sampling approach to explore the impact of intra-specific variation on σ^2^ (see Supplement §5). We found the same pattern of a higher σ^2^ in male autosomal genes compared to female autosomal genes, and a higher σ^2^ of Z genes with no difference between the sexes (Fig. S7), suggesting that the median value does not obscure broad patterns. Together, these results show that gene expression on the Z chromosome evolves ∼25% faster than all autosomes, and among autosomes, male gene expression evolves ∼5% more quickly than female gene expression values, yet there is no sex difference in evolutionary rate of gene expression on the Z.

Looking next at the M:F ratio of gene expression, we found that the degree of sex bias on the Z evolves almost twice as fast as that of autosomes (σ^2^ = 0.00070, vs. 0.00037); this model is a significantly better fit than the model that does not allow for different rates among chromosomes (Table 3). Note that σ^2^-values for this trait are more than an order-of-magnitude lower than for the median gene expression above. This likely reflects the fact that male and female expression are generally highly correlated, so that changes in their relative magnitude (reflected in the M:F ratio) occur more rarely.

### Modelling with distinct rate categories

To allow genes to vary in their evolutionary rate regardless of their chromosomal position or pre-specified hypotheses, we fit a discrete-gamma distribution with 10 rate categories (*k* = 10) to the data from both gene expression (α = β = 1.12887, σ^2^ = 0. 01138) and M:F ratio (α = β = 0.792558, σ^2^ = 0. 00056), and we found significant overlap among rates categories for these two metrics (Fig. 3A; FDR p < 0.05). This indicates that, in general, genes evolve quickly in both their overall expression and their sex ratio (upper right corner Fig. 3A), and genes evolve slowly in both overall expression and sex ratio (lower left corner). However, the median rate of gene expression evolution is not necessarily predictive of pace of sex ratio evolution, as there are many genes in the fastest rate category for M:F ratio that have moderate to low rates for overall expression evolution, and vice-versa (Fig. 3A, top row and right column; see Fig. 3B for example genes).

**Figure 3:**
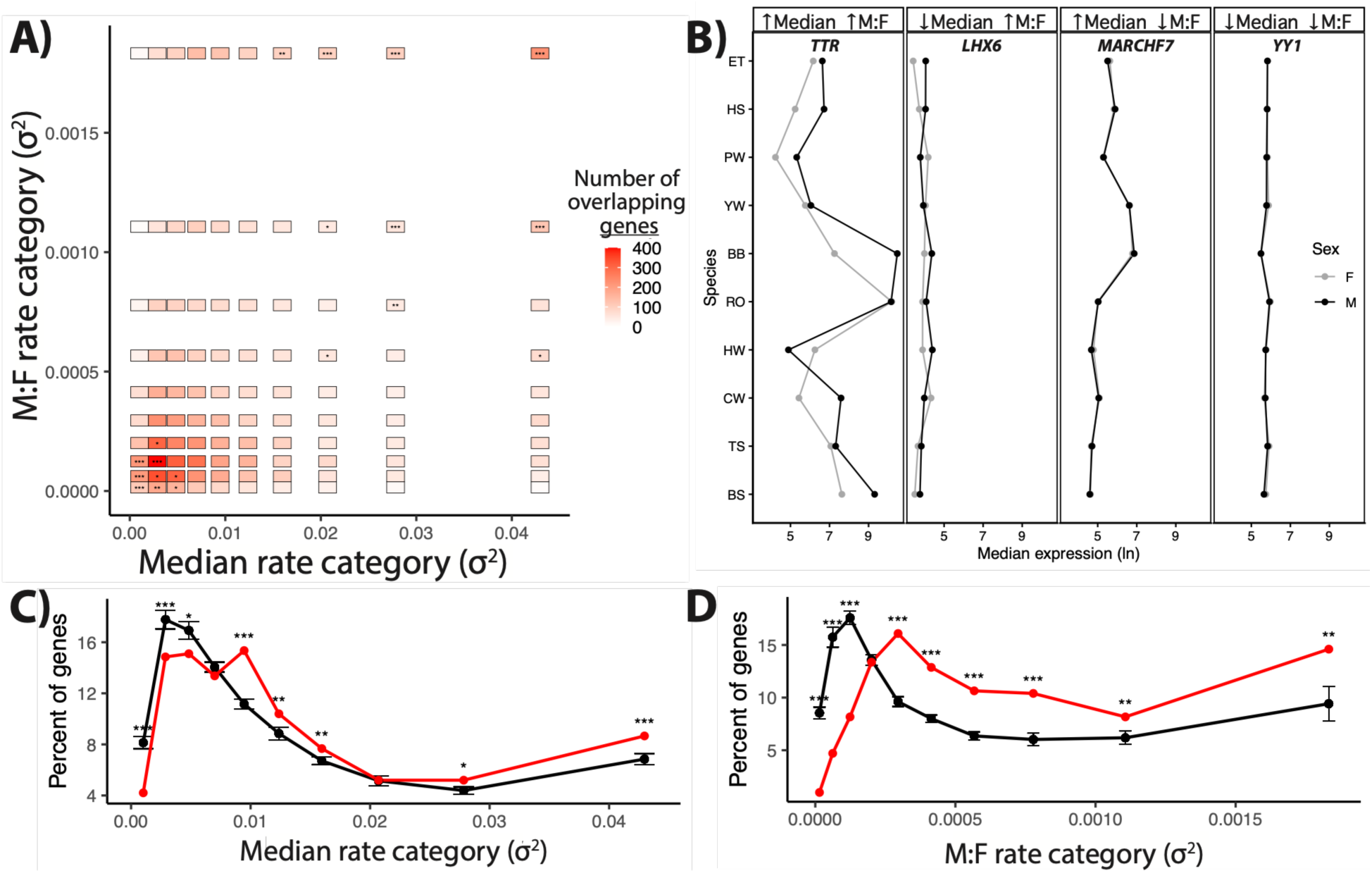
**A)** Comparison of the number of genes in each rate category (σ^2^), based on M:F ratio of expression (y-axis) and median expression (x-axis). Significant enrichment is calculated for every rate-category separately; *p < 0.05, **p < 0.01, ***p < 0.001. **B)** Examples of genes drawn from the corners of panel A, with top panel labels indicating if the gene is assigned to the fastest (up-arrow) or slowest (down-arrow) rate category for median expression and M:F ratio, respectively. For each species (y-axis), the median gene expression (x-axis) is displayed for both males (black) and females (grey). Note that the two lines are highly overlapping in some panels, indicating little difference between males and females. **C)** Percentage of genes assigned to different rate categories (σ^2^) on the Z chromosome (red) and autosomes (black; mean and SE) for median gene expression. **D)** Percentage of genes assigned to different rate categories (σ^2^) on the Z chromosome (red) and autosomes (black; mean and SE) for M:F ratio. Significance is calculated for Z vs. autosomes for every rate-category separately; *p < 0.05, **p < 0.01, ***p < 0.001.

The best-fit gamma distribution shows that most genes evolve slowly, with a long tail of rapidly evolving genes, for both median expression (Fig. 3C) and M:F ratio (Fig. 3D). Among the fastest evolving genes, we found no significantly enriched GO categories for either median gene expression or M:F ratio. For genes with the slowest rate of gene expression evolution, however, there is significant enrichment for GO pathways involved with cell signaling, protein acetylation, and mRNA metabolism (Fig. S6A). For the slowest evolving genes in M:F ratio, there is significant enrichment for GO pathways involved DNA regulation, cell cycle regulation, protein and mRNA processing (Fig. S6B).

Since this multi-rate CAGEE model allows genes to vary in their rate of evolution, without respect to pre-specified groups of genes, we used this model to test again whether genes on the Z are evolving faster than those on autosomes. Consistent with our earlier analyses (Tables 2, 3), the Z chromosome has a higher proportion of genes in the fastest rate categories, and lower proportion in the slowest rate category, compared to the average autosome for both gene expression (Fig. 3C) and M:F ratio (Fig. 3D) (see Supplemental Dataset S3 for full results). Specifically, for gene expression, 8.67% of Z genes were in the fastest rate category (vs. autosome mean = 6.17%, SE = 0.37) and 4.21% of Z genes were in the slowest rate category (vs. autosome mean = 9.15%, SE = 0.38). Similarly, for M:F ratio, 14.60% of Z genes were in the fastest rate category (vs. autosome mean = 5.84%, SE = 0.58) and 0.99% of Z genes were in the slowest rate category (vs. autosome mean = 8.93%, SE = 0.55).

Measures of gene expression can be heteroskedastic, with more lowly expressed genes showing higher variance (even after log-transformation; e.g.(McIntyre et al., 2011)). This relationship could in turn lead genes with generally lower expression levels to be assigned to faster evolutionary rate categories than genes with higher expression levels. This pattern holds in our data, though these effects are not large (see Fig. S5A-C; Supplemental §3). Moreover, we found that the evolutionary rate difference between Z and autosomes is unlikely to be due to this effect because the distribution of Z gene expression levels is not different from that of a similarly sized autosome and evolutionary rates do not differ when comparing the lowest expressed genes between these chromosomes (see Fig. S5D; Supplemental §3).

### Modelling ancestral states of male:female ratio

Focused on this degree of sex bias among Z genes (i.e. M:F ratio, Fig. 2B, red), we found no significant relationship between a gene’s ancestral state (as estimated by CAGEE) and its evolutionary rate (ordinal logistic regression: β =-0.137, SE = 0. 4497, p = 0.761; Fig. 4), in contrast to our second main hypothesis.

**Figure 4:**
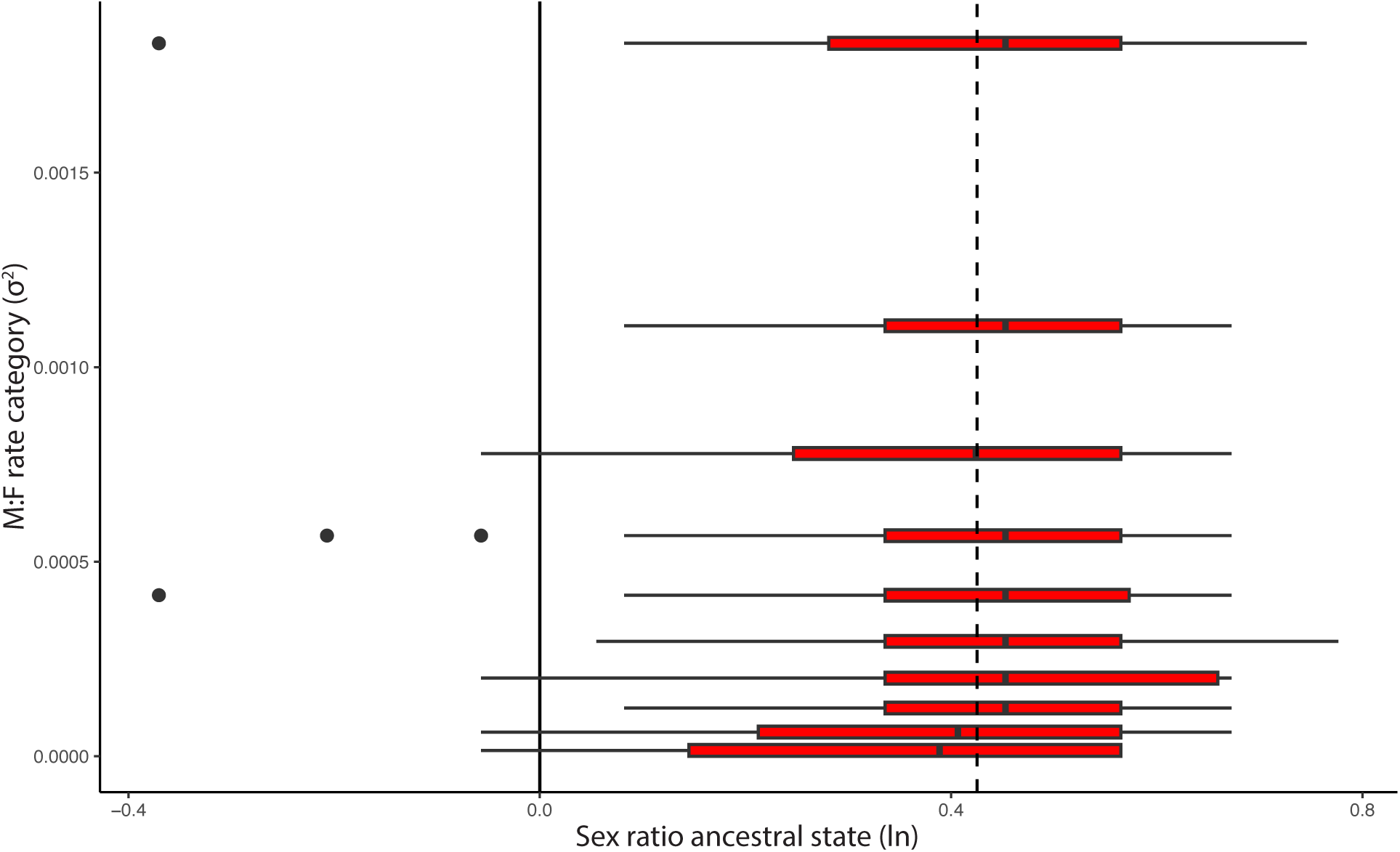
On the Z chromosome, the sex ratio of the ancestral state at the root of the tree (x-axis) is unrelated to the rate of gene expression evolution for M:F ratio (y-axis). Box and whisker plots show distribution of sex ratio ancestral states for Z genes in each M:F σ^2^ category, with outliers shown as points. Dashed vertical line is the average ancestral state for all Z genes.

### Modelling the role of nesting strategy

Regarding nesting ecology and focusing first on median gene expression without considering sex, the best fit model allows for three rates: one for all internal branches along the phylogeny (σ^2^ = 0.00625), another (higher) value for obligate cavity-nesters (σ^2^ = 0.01328), and a third (even higher) value for species that are not obligate cavity-nesters (σ^2^ = 0.01400). Based on likelihood ratios, this three-parameter model is a significantly better fit than alternatives (Table 4). These results indicate that brain gene expression levels evolve ∼5% *slower* along obligate cavity-nesting vs. more flexible-nesting lineages. Analyzing the sexes separately reveals the same pattern (Tables S1, S2), suggesting that cavity-nesting does not differentially affect rates of brain gene expression evolution in one vs. the other sex.

In contrast, we find a different outcome when using the M:F ratio to test our third main hypothesis. Specifically, obligate cavity-nesting species have ∼18% *faster* evolution of M:F ratio compared to flexible-nesting species (σ^2^ = 0.00059 vs. 0.00050; Table 5). Thus, even though median gene expression and sex ratio of expression are not completely independent, these different measures provide different insights on the interplay between sex and transcriptomic evolution.

## DISCUSSION

We developed new features in CAGEE (v1.2, Fig. 1), applied here to test three hypotheses on the evolution of sex-biased brain gene expression in songbirds. In support of our first hypothesis, genes on the Z chromosome exhibit a faster evolutionary rate of median neural gene expression and M:F ratio, compared to autosomal genes. However, among our focal species, there is no strong evidence that gene expression on the Z evolves faster in males than females. Additionally, Z genes with slower evolutionary rates in sex ratio are not largely ancestrally more balanced between the sexes, indicating no relationship between the M:F gene expression ratio and the accumulation of variation in this metric across the phylogeny. This pattern is inconsistent with our second hypothesis, that the relatively balanced expression seen in some Z genes may be a result of historical selection for balance (Mank & Ellegren, 2009). Finally, we find that sex ratios in gene expression evolve more rapidly along songbird lineages that have convergently evolved obligate cavity-nesting, a trait associated with sex-specific changes in selection related to territorial aggression (Lipshutz et al., 2025). Across multiple results, our direct analyses of sex ratio evolution provide different effect sizes from indirect analyses that measure the sexes separately (Table 2 vs. 3; Table 4 vs. 5; Fig. 3C vs. 3D). Together, these findings demonstrate (i) that the sexes need to be studied concurrently to understand how evolution shapes gene expression and (ii) how new features in CAGEE v1.2 can advance the field of comparative transcriptomics.

### Z chromosomes and the evolution of gene expression

All ten focal species displayed similar levels of male-biased expression on the Z (Fig. 2B & Supplemental §1), with Z gene expression averaging 59% higher for males than for females. This unequal gene expression has been documented previously (Itoh et al., 2007; Julien et al., 2012) and is important for the study of sex differences because there is evidence in birds that secondary sexual traits are driven by sex differences in expression of Z genes, even when controlling for differences in gonadal hormones or global methylation (Diddens et al., 2021; Ioannidis et al., 2021; Wade et al., 1999). We extend these findings here, documenting faster gene expression evolution on the Z, coupled with a faster rate of evolution of the sex ratio of expression as well. These higher rates on the Z could be the result of reduced effective population size of the Z compared to autosomes, differences in selection on the two sets of chromosomes, or some combination of these (Dean et al., 2015; Hayes et al., 2020; Ottenburghs, 2022). Furthermore, the higher evolutionary rates we observe do not distinguish among the causes of these differences, which would require future study. However, we do not find evidence for our first main hypothesis of faster gene expression evolution on the *female* Z relative to the male Z (Table 2), as has previously been shown in peripheral tissue of other non-songbirds (Dean et al., 2015). It is therefore possible that, unlike peripheral tissue, neural tissue does not show a pattern of a faster evolutionary rate on the female Z, suggesting the potential for tissue-specific rates of sex ratio gene expression evolution.

With the new unbounded Brownian motion feature introduced here, we now provide a non-mutually exclusive alternative interpretation– that outcomes from direct analyses on the degree of sex bias give different answers than analyses of each sex separately. Modelling gene expression evolution for the sexes separately, though, we found only small differences in the pace of change on autosomes, with male gene expression evolving about 5% faster than that of females (Table 2). This pattern is consistent with a previous analysis in peripheral tissue from Galliform birds, which reported a greater number of evolutionary transitions of male-biased compared to female-biased autosomal genes (Harrison et al., 2015). While our sex-specific analysis of gene expression on the Z vs. autosomes revealed only modest differences, our direct analysis of the degree of sex difference (i.e. M:F ratio) revealed that sex-biased gene expression on the Z evolves almost twice as fast as that of autosomes. Importantly, there is evidence in birds that sex-biased gene expression on the Z may drive tissue-specific sexual differentiation, which is only weakly controlled via sex hormones after gonadal differentiation (Deviatiiarov et al., 2023; Ioannidis et al., 2021; Zhao et al., 2010). Moving forward, it will be critical to apply these direct approaches beyond our focal songbird species and in other tissues, to assess generality. More broadly and beyond the context of sex, the unbounded Brownian motion feature can also apply to model the degree of widely used treatment effects (e.g. response to social challenge *sensu* (Rittschof et al., 2014)), as well as other non-positive continuous data, like those extracted from PCA, WGCNA, or other multi-omics datasets.

### Variable rates of expression evolution among genes

The second new CAGEE feature we developed– variable rates of evolution for different genes on the same branch of the phylogeny – further advances understanding of the evolution of sex-biased gene expression. For one, by assigning genes into ten rate categories without *a priori* hypotheses, we provide a separate inference on the rate of Z vs. the autosomes, mirroring the single rate analyses (Tables 1-3) in that genes with the fastest evolutionary rates are concentrated on the Z chromosome (Fig. 3C,D).

When allowing rates of evolution to vary among genes, it is important to consider that heteroskedasticity could bias rate estimation (Fig. S5A,B). We mitigate this issue through the inherent variance stabilization of CAGEE’s log-transformation and our use of median expression values across samples. Post-hoc analyses indicate that the impact of heteroskedasticity is mild and cannot explain the patterns we observed (See Supplemental §3). Though individual genes could be impacted, heteroskedasticity will have minimal impact when comparing rates across lineages or across the transcriptome, as there is a similar distribution of gene expression across species. In the future, models that directly account for measurement error (see Han et al., 2013) may further mitigate this issue in datasets where heteroskedasticity is particularly severe.

The variable-rate feature also opens doors to asking questions about why some genes evolve more quickly in their degree of sex bias vs. balance Understanding the degree of sex bias is a particularly salient issue in birds due to their lack of global dosage compensation (Fallahshahroudi et al., 2025; Itoh et al., 2007; Julien et al., 2012), noting that our results show marked variation in the degree of sex bias for individual Z genes across different species (Fig. S1). In each of our 10 focal species, we find that some Z-linked genes are expressed relatively equally between females and males (Fig. 2B), consistent with prior work (Mank & Ellegren, 2009; Zimmer et al., 2016). Past comparisons of two or three bird species also provide some evidence that there may be consistency in which Z genes are most balanced between the sexes over the course of evolutionary time (Itoh et al., 2010; Ko et al., 2021; Naurin et al., 2011; Wolf & Bryk, 2011), as would be expected if there is some form of gene-by-gene purifying selection for balance (Mank & Ellegren, 2009). If so, those genes on the Z under selection for a more balanced sex ratio would be conserved, as reflected in both a relatively balanced ancestral state and a slower rate of gene expression evolution in accordance with our second main hypothesis. Contrary to this hypothesis, though, we find no relationship between the rate of gene expression evolution and the degree of sex bias at the ancestral root node of these 10 songbirds (Fig. 4), a finding which implicates relaxed selection as a more likely evolutionary factor than other processes. There is a possibility that this lack of purifying selection manifests in other tissues, life-history stages, or lineages. For example, other bird lineages, but not songbirds, have hypermethylated regions of the Z that may represent the first step towards the evolution of complete dosage compensation (Melamed & Arnold, 2007; Sun et al., 2019; Wright, Zimmer, et al., 2015). Other processes, like transcriptional bursting (Papanicolaou et al., 2025), post-transcriptional compensation (Fallahshahroudi et al., 2025; Lister et al., 2024; Uebbing et al., 2015), gene-specific hormonal effects (van Nas et al., 2009), or variation across developmental time (Mank & Ellegren, 2009) may still operate, and future analyses should further explore how these factors interact with rates of gene expression evolution.

Another non-mutually exclusive hypothesis is that genes with certain functions evolve at different rates. This issue has been explored in depth in the past, demonstrating the slower sequence evolution of essential genes related to the synthesis of DNA, RNA, and proteins across animals (Rancati et al., 2018; Zhan & Boutros, 2016). We approached this question with GO analyses of the genes in the most extreme rate categories, finding no functional enrichment for genes that evolve most quickly– either in terms of their median levels of expression or their degree of sex bias. Focused on genes with the slowest rates of gene expression evolution, however, we do find enrichment of several basic cell functions, including cell cycle, chromatin organization, RNA and protein processing. These findings are consistent with the likely constraints that shape these mission critical pathways, with little wiggle room for gene expression, or its degree of sex bias, to change among species.

Our analyses underscore open questions as to why and how it is that some Z genes exhibit relative balance in their expression despite the fact that males have two copies of the Z and females have just one. Notably in our analyses, genes that evolve most quickly in their degree of sex bias also tend to evolve most quickly in their median expression level. Exceptions occur, with hundreds of genes that evolve quickly in their sex ratio but not their overall expression, and vice versa (Fig. 3A, 3B). Thus, even though these two metrics are related, they can vary semi-independently, including cases in which overall expression evolves slowly but the sex ratio of expression still evolves quickly. We urge future research to explore these relationships across tissues and across different taxa, alongside further optimization of the best number of rate categories for these and other research questions.

### Sex associated gene expression evolution and reproductive ecology

Past research provides key insights on the ecological drivers of sex differences (Gonzalez-Voyer et al., 2022), and we extend this here in relation to obligate cavity-nesting in songbirds, which involves many changes in the phenotype and has evolved convergently across half of the species in our study (Fig. 1A). A previous analysis, using these same transcriptomic data from the ventromedial telencephalon, identified a small set of genes whose expression convergently evolves alongside higher levels of aggression and obligate cavity-nesting (Lipshutz et al., 2025). Here, to test our third main hypothesis, we add new findings linking this element of reproductive ecology with the degree of sex bias in gene expression and the pace of gene expression evolution. Specifically, along obligate cavity-nesting lineages, we find a lower overall rate of gene expression evolution (Table 4), suggesting an ecological constraint on transcriptome evolution. Alternative models that treat the sexes separately are not a significantly better fit (Tables S1, S2), implying that the sex-related gene expression does not evolve differently alongside the convergent evolution of obligate cavity-nesting. Critically, though, with the new ratio-feature in CAGEE v1.2, we see that the M:F expression evolves significantly *faster* along obligate-cavity nesting lineages (Table 5), an insight we would have missed from studying the sexes independently or without regard to the degree of sex bias. These results suggest that, in association with a key component of reproductive ecology, brain gene expression is evolving more slowly in absolute terms, but the expression relationship between males and females is more volatile, with shifts that are more marked in one sex versus the other. While we are unable to determine causality, we speculate that this ecologically-associated acceleration of sex bias evolution relates to the need for different mechanisms to drive convergent behavior across the sexes (sensu: (De Vries, 2004; Ko et al., 2021). More broadly, these results speak to an emerging view that research on sex and sex bias should consider the degree of sex differences (McCarthy et al., 2012; Smiley et al., 2024) as we seek to understand the interplay between sex and evolutionary processes.

## ACKNOWLEDGMENTS

For discussion and advice, we thank members of the Rosvall and Hahn labs, as well as three anonymous reviewers. For support in data collection and sequencing, we thank AB Bentz, AM Buechlein, TA Empson, EM George, ME Hauber, DB Rusch, WM Schelsky, AM Turner, SE Wolf, MJ Woodruff, and the IU Center for Genomics and Bioinformatics.

## DATA AVAILABILITY STATEMENT

All data are available in the main text and/or supplementary materials, with scripts available on Zenodo (Miller-Crews, 2026). The CAGEE software is available at https://github.com/hahnlab/CAGEE.

## BENEFITS GENERATED

Benefits of this research come from the public sharing of our data and results, discussed above.

## AUTHOR CONTRIBUTIONS

SEL, IMC, MWH, and KAR conceived of the study; SEL led the sample collection and sequencing. IMC analyzed and interpreted the data, supported by BF and JB. BF, JB, and MWH developed computational tools. IMC, MWH, and KAR wrote the paper, and all authors contributed to revisions. The manuscript reflects the contributions and ideas of all authors.

