## Supplemental for "How sex shapes transcriptome evolution in the songbird brain"

AFFILIATIONS
1: Department of Biology, Indiana University, Bloomington, IN, USA
2: Department of Biology, Duke University, Durham, NC, USA
3: University Information Technology Services, Indiana University, Bloomington, IN, USA
4: Department of Mathematics, Western University, London, ON, Canada
5: Department of Computer Science, Indiana University, Bloomington, IN, USA

This file includes:

Legends for Datasets S1 to S3
Supplementary text §1 to §5
Figures S1 to S7
Tables S1 to S2
References for supplementary text

### Dataset Legends

**Dataset S1 (separate file).** Normalized gene expression across samples from all 10 songbird species.

**Dataset S2 (separate file).** Differentially expressed genes between males and females for each species. Chromosomal information (chromosome_name) is provided along with DEG analysis results including Log2FoldChange values (LogFC), FDR adjusted p-values (padj), and species.

**Dataset S3 (separate file).** Evolutionary rate assignment for individual genes across 10 species songbird phylogeny. Chromosomal information is provided along with species median gene expression evolutionary rate (median_sigma), and M:F ratio evolutionary rate (M:F_sigma).

*§1. Differential expression analysis of sex bias per species*

As a window into the scope of sex-biased gene expression in our dataset, we used limma-voom (Law et al., 2014) to identify genes that are differentially expressed between female and males within each species (hereafter, 'DEGs'). We operationalized 'male-biased' and 'female-biased' DEGs as significantly higher in males or females, respectively (adjusted p-value $< 0.05$). On average, 248 genes (SD = 81.8) were male-biased and 56 genes (SD = 53.7) were female-biased, per species (adjusted $p < 0.05$) (Fig. S1A). Among the ~10k orthologous genes in our dataset, this amounts to only 1.85% of genes expressed in the ventromedial telencephalon being sex-biased (range from 1.01% to 3.56% of genes, depending on the species). Sex differences in gene expression appear to be mainly species-specific outside of male-bias on the Z chromosomes (Fig. S1B,C,D). Importantly, no autosomal genes were consistently sex-biased across all species, with only 33 autosomal genes having a sex-biased pattern in more than one species and only one autosomal gene having a sex-biased pattern in five species (Fig. S1B).

A majority of the male-biased DEGs occurred on the Z chromosome, with on average 190 out of the 404 Z-mapped genes being male-biased DEGs (SD = 41.4) (Fig. S1C). Very few female-biased genes occurred on the Z, with on average 2.2 out of 404 Z-mapped genes being female-biased DEGs (SD = 0.8) (Fig. S1D).

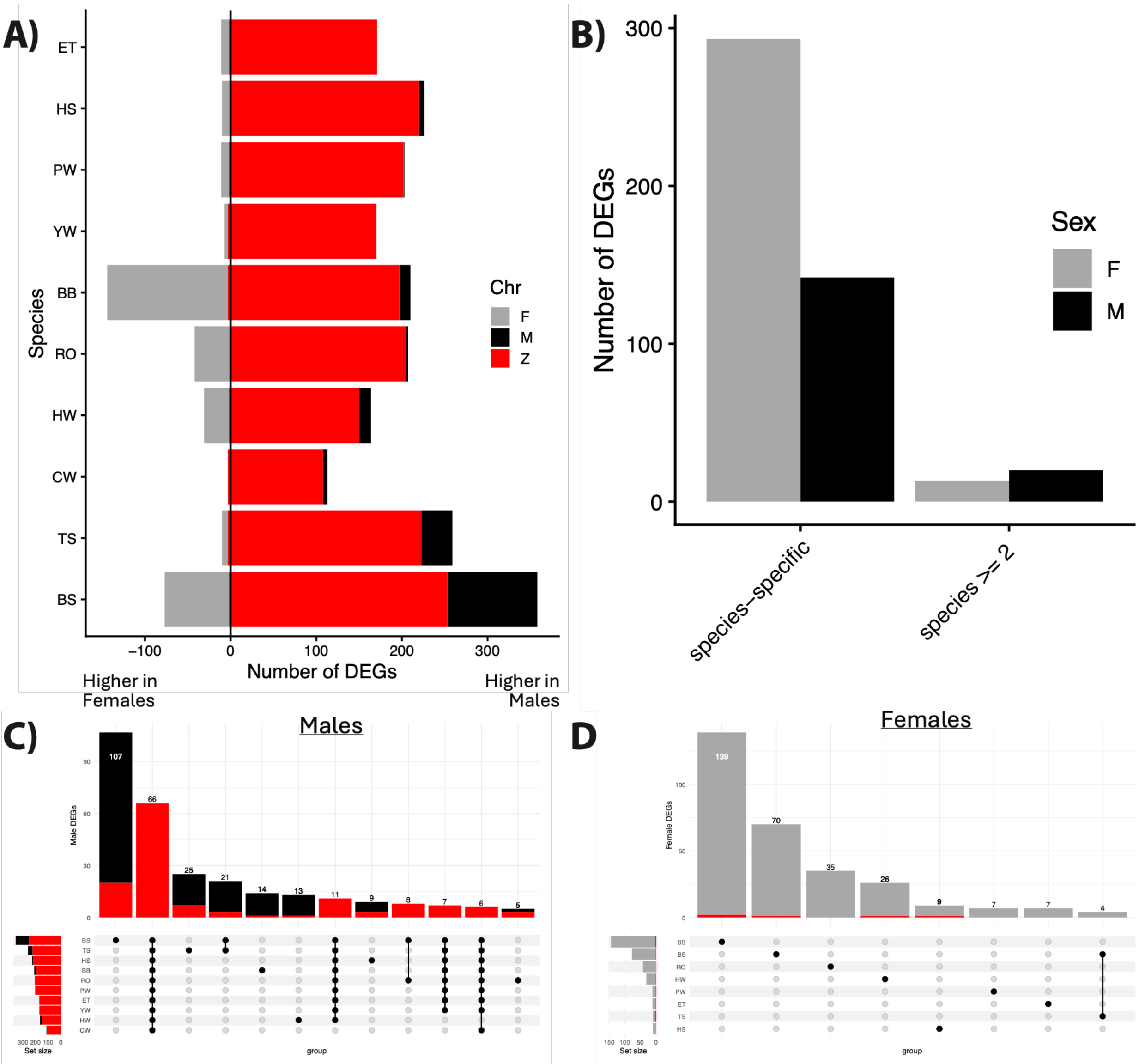


*Figure S1: Sex-associated differential gene expression across 10 songbird species.*

**A)** Bar plot shows, per species, the number of autosomal DEGs that are male-biased (right; black) or female-biased (left; grey), and number of DEGs on the Z chromosome (red).

**B)** The number of DEGs on autosomes that are male-biased (black) or female-biased (grey) only in a single species ('species-specific') or are consistently biased across multiple species ('species >= 2').

Upset plots show the overlap of **C)** male-biased DEGs among species and **D)** female-biased DEGs among species, with Z genes marked in red.

To look beyond the number of sex-biased genes and their (limited) overlap across species (Fig S1B,C,D), we used CAGEE v1.2 to identify whether genes that evolved in the degree of M:F ratio did so in the same direction across all external branches. Because of uncertainty in ancestral state reconstruction, CAGEE denotes changes outside the possible range of probable ancestral states as a "credible change" (Bertram et al., 2023). For each gene, we count the number of credible changes across branches and the direction of those changes.

CAGEE 1.2 estimated that 5,308 of 10,672 genes showed a credible change in M:F ratio on at least one branch (Fig. S2A). This means that roughly half of expressed genes became either more male-biased or more female-biased during their evolutionary history. Importantly, this does not indicate that the genes became sex-biased or not – only that they shifted in their relative degree of sex-bias. Of those, only 3,244 genes have any changes on two or more lineages, and only 266 genes are consistently in the direction of change across multiple changes (Fig. S2B). In short, there were few genes that changed in the exact same manner across all species, consistent with the classic DEG approach (above) (see also (Harrison et al., 2015)), which reveals little concordance in which genes are sex-biased across species.

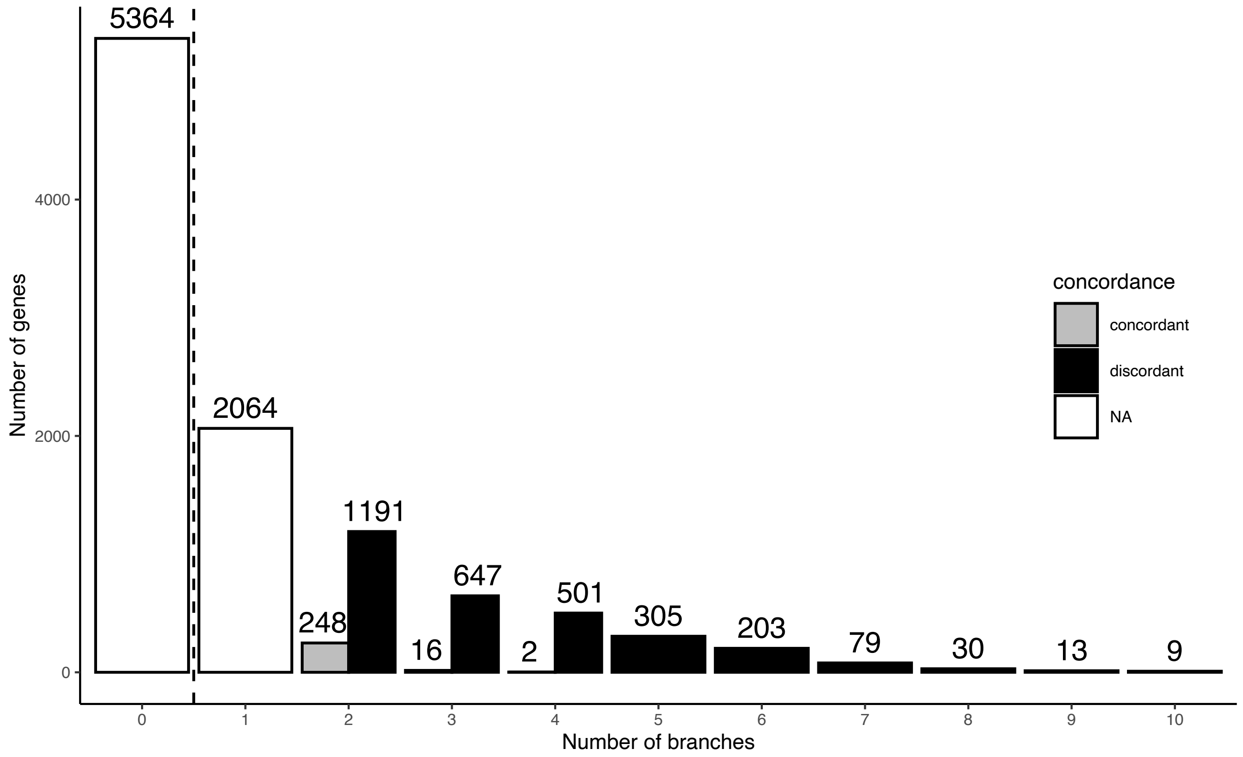


*Figure S2: Credible changes in M:F ratio across 10 songbird species.*
For genes with credible changes on 2 or more external branches, we note whether those changes are in the same direction (concordant, gray) or in opposing directions (discordant, black). For example, there are 1439 genes that show a credible change in their sex ratio on just two branches, yet most genes (1191) change discordantly, becoming more male-biased in one lineage while becoming more female-biased on another lineage. There are only 9 genes that show credible changes across all 10 branches, and these do not change concordantly.

*§2. Simulation of variable evolutionary rate among genes.*

To test the new feature allowing rate variation among genes, four sets of 100 genes with different values for $\sigma^2$ were simulated in CAGEE. This process results in a single combined dataset with four sets of genes from four different simulated values of $\sigma^2$. We created four of these datasets, for a range of both large and small differences in $\sigma^2$, as well as for both gene expression and M:F expression ratio. We used these combined datasets to test whether the gamma-distributed rate feature could both successfully separate genes into their four separate categories and estimate accurate values of $\sigma^2$ for each category. Using the simulated datasets, we show that genes are assigned to rate categories as expected and that the $\sigma^2$ of those categories closely matches the known simulated $\sigma^2$ for those genes (Fig. S3).

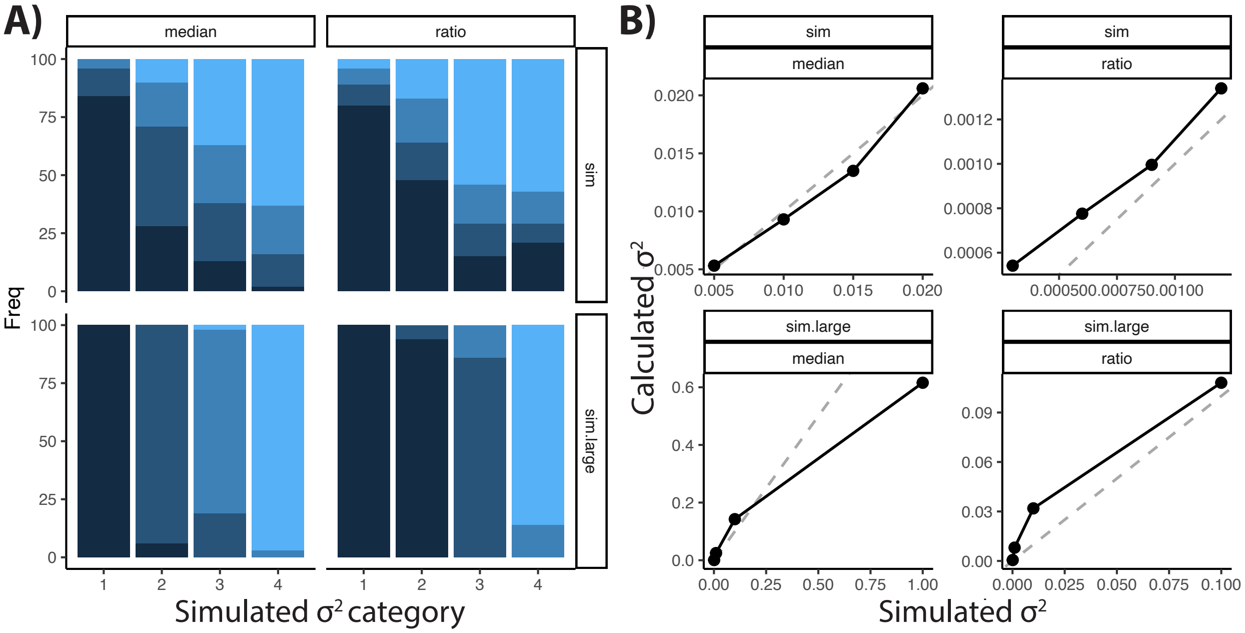


*Figure S3: Simulation results testing new rate category feature in CAGEE v1.2*
**A)** Number of genes out of 100 genes from each simulated rate category (1 = slowest, 4 = fastest) for each of the four datasets of simulated genes assigned to each relative rate category (dark blue = slowest, light blue = fastest). **B)** Relationship between simulated $\sigma^2$ and calculated $\sigma^2$ for each of the four datasets of simulated genes (grey, dashed line = 1:1 line). Left side is simulated gene expression data ('median') and right side is simulated gene expression ratio data ('ratio'). Top row and bottom row are using small changes ('sim') and large changes ('sim.large) in $\sigma^2$ across categories, respectively.

Additionally, to test the consistency of this new rate-category feature, we analyzed data across a range of categories (*k* =2, 4, 6, 8, 10) for both species (median) gene expression and M:F ratio (Fig. S4). We found that $\sigma^2$ changes slightly across *k* categories, with an increased rate at low values of *k* before plateauing as *k* increases, for both median and M:F (Fig. S4A). This pattern mirrors previous results using similar models and is a known feature of increasing *k* (Mendes et al., 2020). As increasing values of *k* will produce more rate categories, we also wanted to test the degree of consistency in both gene membership and assigned $\sigma^2$, for both median and M:F (Fig. S4) as *k* values increase. As expected, we found that higher *k* improved resolution, as rate categories are consistently split into two smaller rate categories. However, these splits are not always equal, as is apparent in the genes assigned to the fastest rate categories being assigned consistently to the fastest rate category across values of *k* (Fig. S4). While it is not possible to pick an optimal value for *k*, we targeted *k* = 10 to increase resolution while maintaining on average 1,000 genes per rate category.

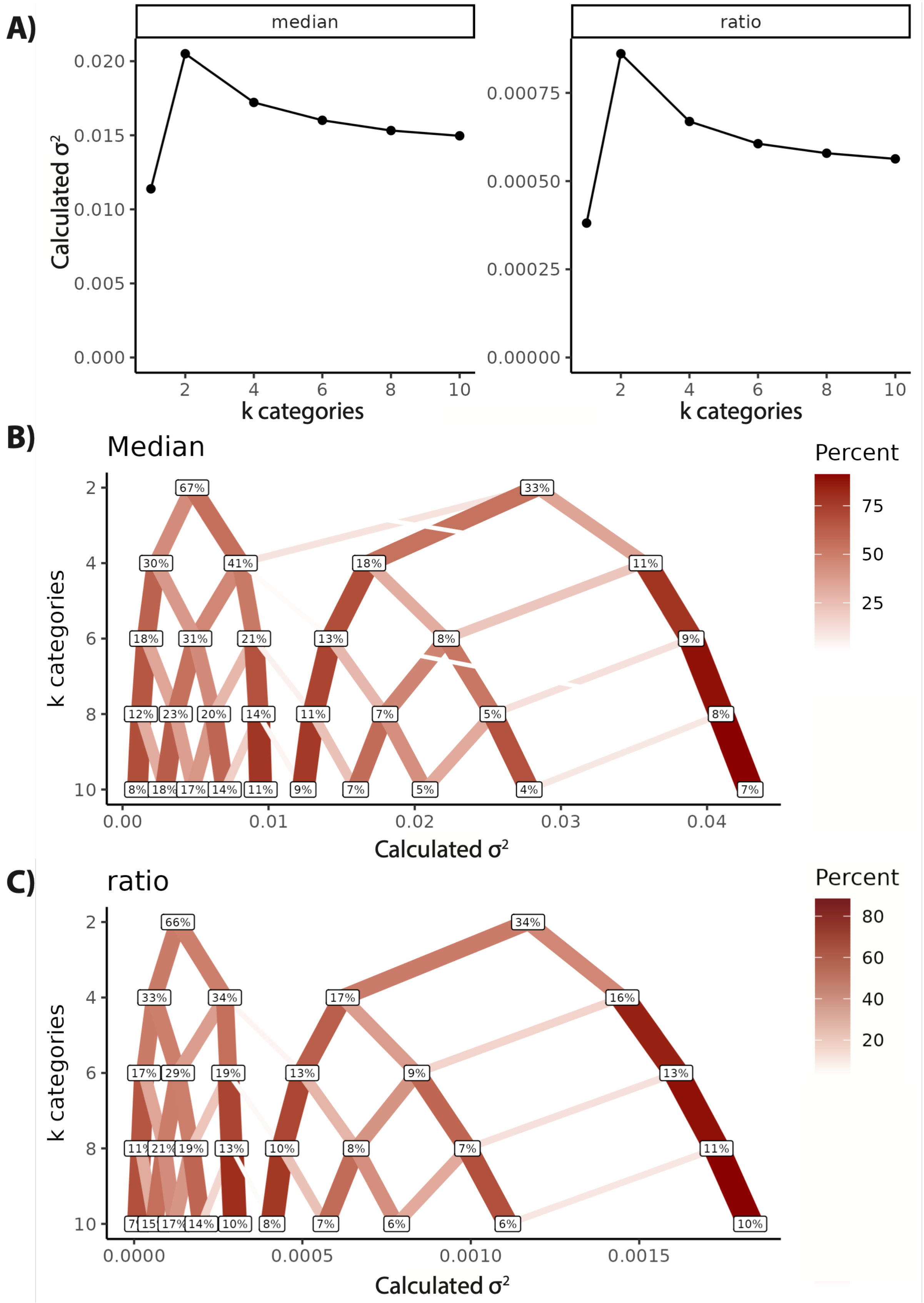


Figure S4: *Results with varying numbers of rate categories*

**A)** Calculated $\sigma^2$ across values of k (k = 1, 2, 4, 6, 8, 10) for both median (left) and M:F (right) gene expression. For median gene expression (**B**) and M:F (**C**), the percentage of genes in each rate category (label) across values of k (y-axis) and the assigned $\sigma^2$ for that rate category (x-axis). Edges connecting the labels (red lines) show the percentage of genes (line color and width) from the rate category of the lower k value (above) that are assigned to the rate category of the next highest k value (below).

*§3. Effect of gene expression level on evolutionary rates among genes.*

Due to the heteroskedastic nature of gene expression data, which can lead to a relationship between gene expression level and measured variance, we analyzed the relationship between expression level and assigned rate category. In our analysis, the normalized gene expression data encounters two steps to help mitigate the impacts of measurement error. First, we take the median expression within a species, which should reduce outlier bias. Second, the first step in CAGEE is a log-transformation of the expression data, after adding an initial offset value, which serves to stabilize variance. To understand what impact this combination of steps may have on limiting heteroskedasticity, we show that there is a limited relationship between expression level and variance, excepting slight increase in variance for the lowest expressed genes (Fig. S5A). This increased variance appears to be most prevalent for the few genes with an average normalized expression across all species <10 (Fig. S5B), which could be due to higher expression in one or a few species.

To test if this increased variance leads to higher calculated evolutionary rates, we compared evolutionary rate category assignments from species median gene expression to the predicted ancestral state gene expression (Fig. S5C). We found that genes assigned to faster evolutionary rate categories tended to have slightly lower expression levels. To quantify this effect, we compared the assigned evolutionary rate between with lower average expression across species to those with higher average expression across species using a normalized expression cutoff of ~150 (ln(exp) < 5 vs ln(exp) >= 5). We found that these lower expression genes indeed have significantly higher assigned expression rates (t(7324.1) = 23.76, p-value < 2.2e-16).

To ensure that the higher evolutionary rates found on the Z are not related to lower expression levels of the Z compared to autosomes, we compared the relationship between expression level and evolutionary rate between the Z and a similar-sized autosome, chromosome 4. To test if expression level on the Z was lower than expected, we compared the proportion of genes below the median expression level of the entire transcriptome on the Z and chromosome 4 (Fig. S5D). We separated out males and females, as female Z expression level is impacted by a lack of dosage compensation. We found no difference in the proportion of genes below the median expression level of the entire transcriptome between the male Z and either the male or female chromosome 4 (Male Z = 46.3%; Male chromosome 4 = 46.7%; Female chromosome 4 = 45.7%; Female Z = 63.1%). To test if lower expressed Z genes had higher evolutionary rates than lower expressed genes on chromosome 4, we compared evolutionary rates of the bottom 50% expressed genes on the Z to the bottom 50% expressed genes on chromosome 4 and found no significant difference in evolutionary rate (t(426.62) = -1.61; p-value = 0.246). Likewise, we found no significant difference between the top 50% on either chromosome (t(438) = 0.176; p-value = 0.861).

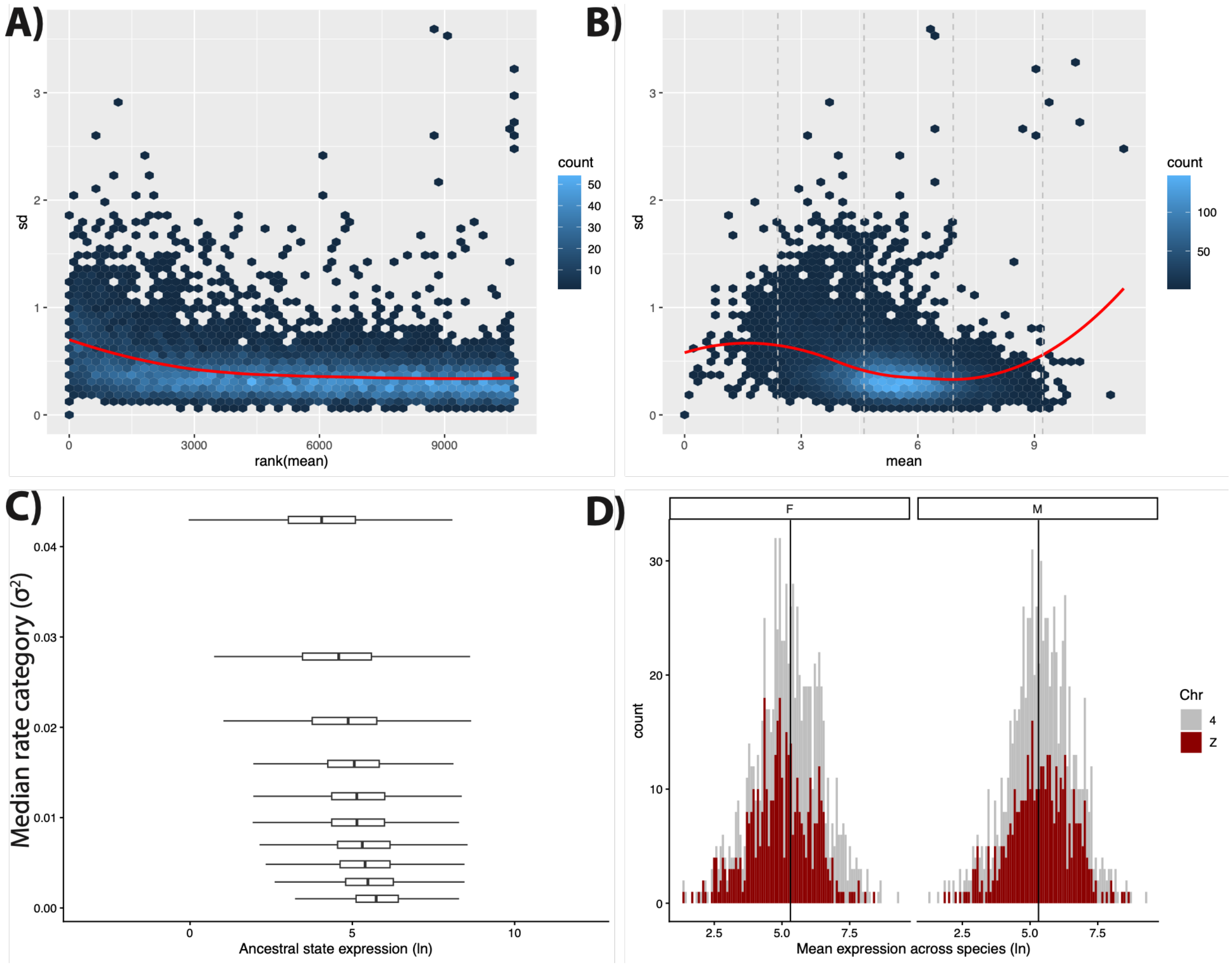


Figure S5: *Impact of gene expression level on estimated evolutionary rate*
**A)** Dispersion plot of the natural log-transformed (ln+1) median of normalized gene expression counts within each species (i.e. each gene has one value per species). Genes are ranked from lowest average expression to highest average expression (x-axis) compared to standard deviation (y-axis). Color indicates the number of genes in each hex-cell and the red line indicates line of best fit. B) Same dispersion plot as in A, with gene expression values themselves instead of log-counts. Vertical, grey dashed lines indicate non-transformed expression levels corresponding to 10, 100, 1,000, and 10,000 respectively. C) Comparison between median gene expression rate category (σ2; y-axis) and the predicted ancestral state gene expression at the root node (x-axis). D) Histogram distributions of average gene expression levels across species median expression (x-axis) for females (left) and males (right) between the Z chromosome (red) and the similarly sized autosomal chromosome 4 (grey). Together, these plots demonstrate that low expression has minimal impact on increasing estimated evolutionary rate. Additionally, the Z chromosome is not impacted by this effect, as the Z has a similar distribution of gene expression levels to autosomes.

### *§4. Functional enrichment in evolutionary rate categories*

To ask if faster- or slower-evolving genes were functionally enriched, we used Ensembl to assign gene ontology (GO) terms for Zebra finch and AnnotationForge (Carlson & Pagès, 2024) to generate our own GO database, containing 8016 genes with GO terms. To calculate GO enrichment, we used enrichGO from the package clusterprofiler (Wu et al., 2021) with a FDR corrected adjusted p-value cutoff of 0.1, minGSSize = 10, and maxGSSize = 500. To graph our results faster- or slower-evolving genes, we used pairwise_termsim to get a hierarchical ordering of GO terms and treeplot for plotting (Fig. S6), both from the package enrichplot (G, 2025).

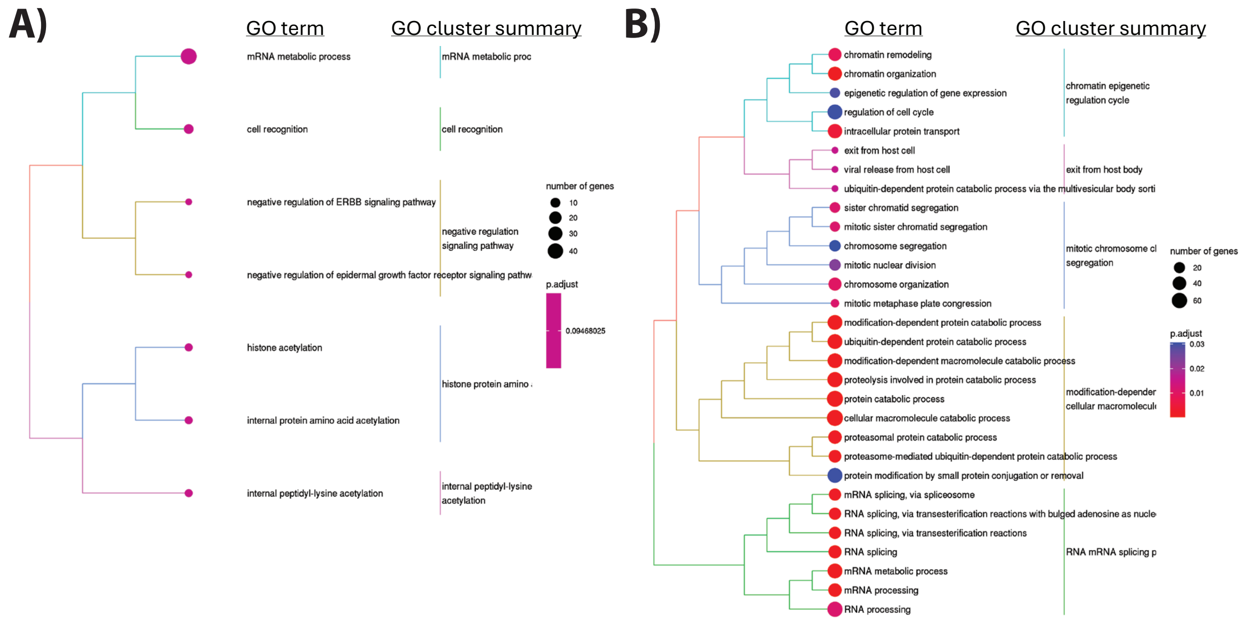


*Figure S6: GO analysis and evolutionary rate*
Genes in the slowest evolutionary rate category for both **A)** gene expression and **B)** sex ratio gene expression show enrichment of gene ontology categories related to basic cell processes. Significantly enriched GO terms are hierarchically clustered (tree) with node tips (circle) representing the adjusted p-value (color) and number of genes in that category (size), next to the name of that GO term. Closely related GO terms share the same branch colors and are summarized by semantic similarity ('GO cluster summary').

*§5. Testing the impact of intra-species variation on* $\sigma^2$

To test the impacts of intra-specific variation on differences in $\sigma^2$, we compared the CAGEE results obtained using median gene expression to those obtained using a randomly subsampled individual. Towards this end, for each of 100 datasets, we randomly sampled one individual male and one individual female as representatives from each species. For each of these datasets, we used CAGEE to calculate male- and female-specific $\sigma^2$ for Z and autosomal genes across all 10 species. We chose the full four-parameter model for this sub-sampling validation to clearly see if there are any sex differences in rates that were obscured from our main analyses that used group median values (Table 2) which reduces intraspecific noise. Using these 100 models, we calculated the 95% confidence interval for $\sigma^2$ of each of the model parameters (Fig. S7). We found the same pattern of results as with the median gene expression (Table 2): Z genes have a significantly higher $\sigma^2$ compared to autosomal genes, yet no difference between the sexes, while male autosomal genes have a significantly higher $\sigma^2$ than female autosomal genes. Of note, the $\sigma^2$ values calculated by this resampling approach are higher than the $\sigma^2$ values calculated using species median gene expression, which is expected as the median gene expression measure may mask noise from individual samples.

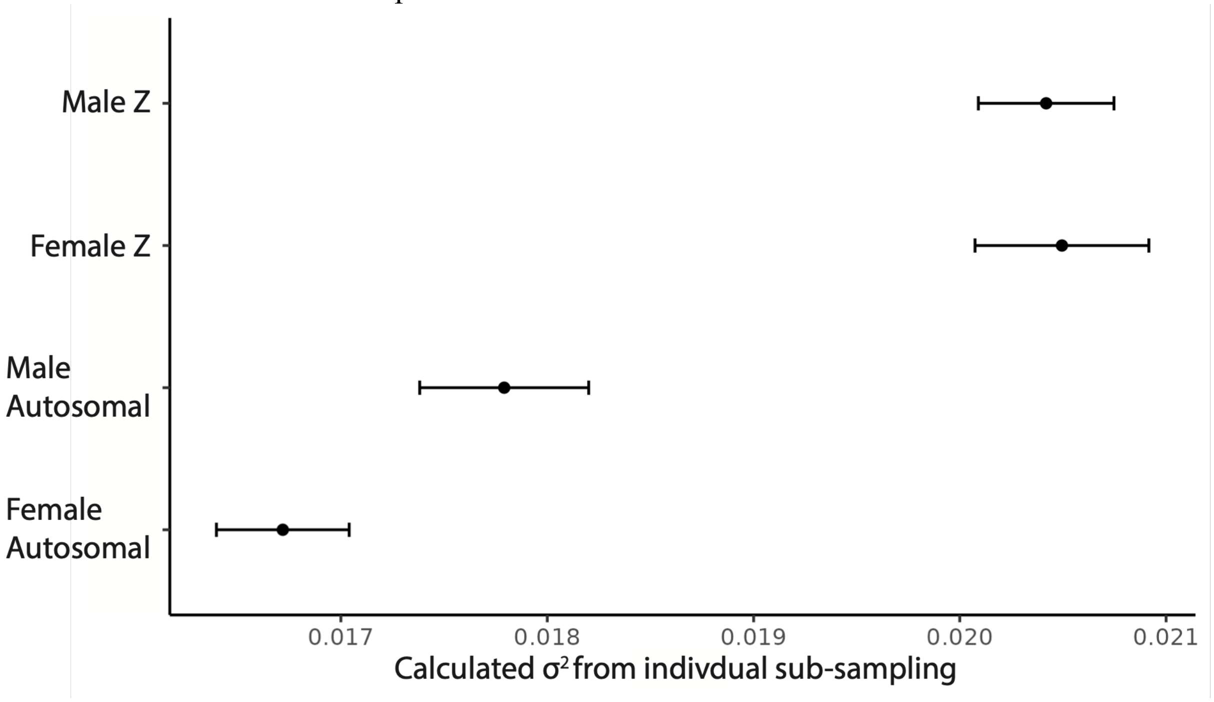


*Figure S7: Sex differences in* $\sigma^2$ *from individual-level resampling*
The 95% confidence intervals and median $\sigma^2$ for male Z-, female Z-, male autosomal-, and female autosomal-genes from 100 models each built from randomly sub-sampling a representative individuals' gene expression from each sex of the 10 songbird species.

**Table S1. Male gene expression in relation to nesting phenotype**

| Model | -lnl | rate | $\sigma^2$ |
|---|---|---|---|
| Males | 340780 | Males | 0.01220 |
| Males: Internal vs External | 336583 | Internal | 0.00626 |
| | | External | 0.01484 |
| **Males: Nesting Phenotype** | **339405** | **Internal** | **0.00619** |
| | | **Obligate** | **0.01461** |
| | | **Flexible** | **0.01518** |

**Table S2. Female gene expression in relation to nesting phenotype**

| Model | -lnl | rate | $\sigma^2$ |
|---|---|---|---|
| Females | 339439 | Females | 0.01189 |
| Females: Internal vs External | 338053 | Internal | 0.00619 |
| | | External | 0.01430 |
| **Females: Nesting Phenotype** | **338045** | **Internal** | **0.00613** |
| | | **Obligate** | **0.01392** |
| | | **Flexible** | **0.01476** |
